# DNA metabarcoding samples from nest soil and beaks outperform samples from talons and faeces when identifying diversity of white-tailed eagle diet in South Greenland

**DOI:** 10.64898/2026.10.14.759070

**Authors:** Mael Perroud, Kay Lucek, Knud Falk, Nicholas Per Huffeldt

**Affiliations:** Biodiversity Genomics Laboratory, Institute of Biology, University of Neuchâtel, Switzerland; Department of Birds and Mammals, Greenland Institute of Natural Resources, 3900 Nuuk, Greenland; Independent Researcher, vandrefalk.dk, Stockholm, Sweden

**Keywords:** *Haliaeetus albicilla*, raptor, diet analyses, foodweb, Arctic biodiversity, plastic pollution

## Abstract

DNA metabarcoding is an emerging approach to study diets across a broad range of species but requires careful optimisation of the sample collection method and substrate used. The suitability of different sample types to identify the diet of the target species remains under investigated. Here, we address these knowledge gaps by collecting four different sample types - faeces, beak swabs, talon swabs, and soil samples - during nest visits of white-tailed eagles (*Haliaeetus albicilla*) during the 2025 breeding season in South Greenland. We further investigated whether some eagle dietary taxa could act as vectors for microplastic pollution. Our analyses revealed a broad diet, with 25 dietary taxa for white-tailed eagles. Lumpfish (*Cyclopterus lumpus*) was the most frequently detected dietary taxon in the diet, which was composed of 75 % fish, 19 % birds, and 7 % mammals. Our findings indicate that diets varied strongly between nesting sites. Although each sample type was efficient in capturing diet, soil and beak swabs were most promising for eagles and gave comparable results. On the other hand, faeces provided a distinct dietary profile with a reduced detection of fish, while requiring an increased sample size, and talon swabs missed rare food items. Finally, we recovered microplastic contamination in 33 % of our samples, but no dietary taxon was associated with microplastic pollution. Taken together our results underline the overlooked value of soil from nests to provide dietary information and highlight the necessity to deepen our knowledge of Arctic food webs and their microplastic contamination.

## Introduction

Studying the trophic interactions and diet of animals is key to understanding their ecology because diet reflects how individuals interact with and respond to their environment (Abrams 2000). Diet can influence species morphology, behaviour, survival, habitat use, distribution and the stability of an ecosystem (Pianka 1969; Abrams 2000; Prigioni et al., 2008). Diet is a fundamental link between organisms and their environment that provides not only nutrients for survival and reproduction but also a basis for interspecific interactions (Prigioni et al., 2008). Traditional methods to study diet rely mainly on direct observation (Wille and Kampp 1983), visual assessments of faeces and stomach contents (Jordan et al., 2005; Pande et al., 2018), kill site examination (Ekblad et al., 2020), or targeted DNA barcoding of pieces of food remains (Méheust et al., 2015). However, those traditional methods have multiple limitations, including difficulties of identifying partially digested food items or limited ability to observe nest sites, and can introduce biases (Nielsen et al., 2018). For instance, visual assessment of faeces from birds, could result in under-represented soft-body food items, like fish, in the diet over items with harder tissues, like mammals, because of differential digestion (Jordan et al., 2005).

Genomic approaches such as DNA metabarcoding (Hacker et al., 2021; Yuan et al., 2024) now permit the identification of dietary taxa from various substrates, including faeces (Liu et al., 2023; Ingvaldsen et al., 2024; Yuan et al., 2024; de Leeuw et al., 2024; O’Hanlon et al., 2025), pellets (Hacker et al., 2021), soil from nests (Good et al., 2024), and swabs of internal or external surfaces (Bourbour et al., 2019; Paprocki et al., 2024; Evans et al., 2025). Faeces are commonly used in birds as their sampling is opportunistic and non-invasive (Yuan et al., 2024; O’Hanlon et al., 2025), but their age can impact the results (McInnes et al., 2017; Cavallo et al., 2018; de Leeuw et al., 2024). Nest soil samples can also be used, but dietary DNA is often scarce (Good et al., 2024). Soil samples can, however, provide additional dietary information as soil captures dietary DNA over longer time scales (Good et al., 2024). Beak and talon swabs also prove effective in dietary identification, but involve direct bird handling (Bourbour et al., 2019; Paprocki et al., 2024). By using a metabarcoding approach on soil, faecal, beak, and talon samples, we assessed the utility of these sample types to overcome limitations of traditional methods by assessing the diet of the white-tailed eagle (*Haliaeetus albicilla*) in Greenland. A better understanding of their diet is relevant for ecological and conservation purposes, as raptors occupy a high trophic position and are an indicator of ecosystem health (Bondì et al., 2016; Castagna et al., 2024). White-tailed eagles also have important roles as ecosystem services providers and as conservation umbrella species (Donázar et al., 2016), but remain suspects of livestock depredation in different parts of the world, including Greenland (Krone et al., 2004; Jaspers et al., 2013).

The white-tailed eagle is a large raptor occurring across Eurasia and the North Atlantic islands, including Iceland and Greenland. The species has become regionally extinct due to lead poisoning (O’Rourke 2014; Nadjafzadeh et al., 2016b), pesticide intoxication, and human persecution (Ekblad et al., 2020). Following conservation measures (O’Rourke 2014), its populations rebounded, but the species is still of conservation importance in several countries (Balotari-Chiebao et al., 2016). The Greenlandic population represents the westernmost population and is genetically isolated from other populations (Hailer et al., 2007; Hansen et al., 2022; Hansen et al., 2023). The species is an opportunistic scavenger and a generalist predator that chooses prey based on their abundance, size, and anti-predator activity to minimise hunting efforts (Nadjafzadeh et al., 2016a). It hunts mainly fish, various bird species, and to a lesser extent mammals (Egede 1770; Wille 1979; Wille and Kampp 1983; Whitfield et al., 2013; Ekblad et al., 2020; Reid et al., 2023; Yuan et al., 2024), but stable isotopes suggest that their diet varies among populations (Nadjafzadeh et al., 2016b). White-tailed eagles may still be persecuted, *i.e.,* shot by humans, as several conflicts exist, *i.e.,* with harvesting common eiders and their down in Iceland and around the Baltic Sea (Ekblad et al., 2020), reindeer herding in Northern Finland (Ekblad et al., 2020), and sheep farming in Scotland, Ireland (O’Rourke 2014), and Greenland (Krone et al., 2004; Jaspers et al., 2013). Beyond trophic interactions, diet can also be used as an indicator of exposure to pollution (Eva et al., 2005). Environmental contaminants such as macro- (>5mm) and microplastics (<5mm) become available in the environment from human sources. Because they degrade very slowly, they accumulate in ecosystems and along the food chain (Mansfield et al., 2024). They could impact top predators, such as raptors of terrestrial and marine ecosystems, as their prey accumulate micro- or macroplastics (Le Guen et al., 2020; Deoniziak et al., 2022; Nessi et al., 2022; Mendez-Sanhueza et al., 2023). Even in remote areas of Greenland, microplastics are present in the ocean, where little auks (*Alle alle*) have been reported to ingest plastics through food (Amélineau et al., 2016). However, whether and to which degree they are present in top predators, like the white-tailed eagle, in Greenland remains unknown.

Here, we implemented a DNA metabarcoding approach to study the diet of the white-tailed eagle in South Greenland during the 2025 breeding season. By comparing four different sample types – faeces, soil from nests, beak swabs, and talon swabs – we aimed to evaluate the utility of different sampling strategies to efficiently characterise the diet of our focal species. We further evaluated the occurrence of microplastics ingested by eagles and tested for an association with their diet. We predicted that the diet of eagles would be broad and mainly composed of fish (Kampp and Wille 1979; Wille and Kampp 1983; Ekblad et al., 2020). We also predicted that swabbing beaks would be the most informative sample type because it likely detects food items longer after their ingestion than faeces (Paprocki et al., 2024). We further predicted seabird species to be the main driver of plastic contamination in our white-tailed eagle population (Provencher et al., 2014; Amélineau et al., 2016).

## Method

### Study area

During the 2025 breeding season, we collected samples in the fjords near the settlements of Narsarsuaq, Qaqortoq, and Narsaq in South-West Greenland (Fig. 1), representing a core breeding area of the white-tailed eagle in Greenland (Hansen 1979; Kampp and Wille 1990) and an area in which the white-tailed eagle was previously studied intensively (Christensen 1979; Ferdinand 1979; Kampp and Wille 1979; Wille and Kampp 1983; Kampp and Wille 1997). The climate ranges from low to sub-Arctic in the fjords (CAFF 2013). We visited seven different white-tailed eagle nests (Fig. 1; Appendix A, Table S1), of which we visited three nests twice. We sampled 11 individual chicks, of which we sampled six chicks twice.

**Figure 1:**
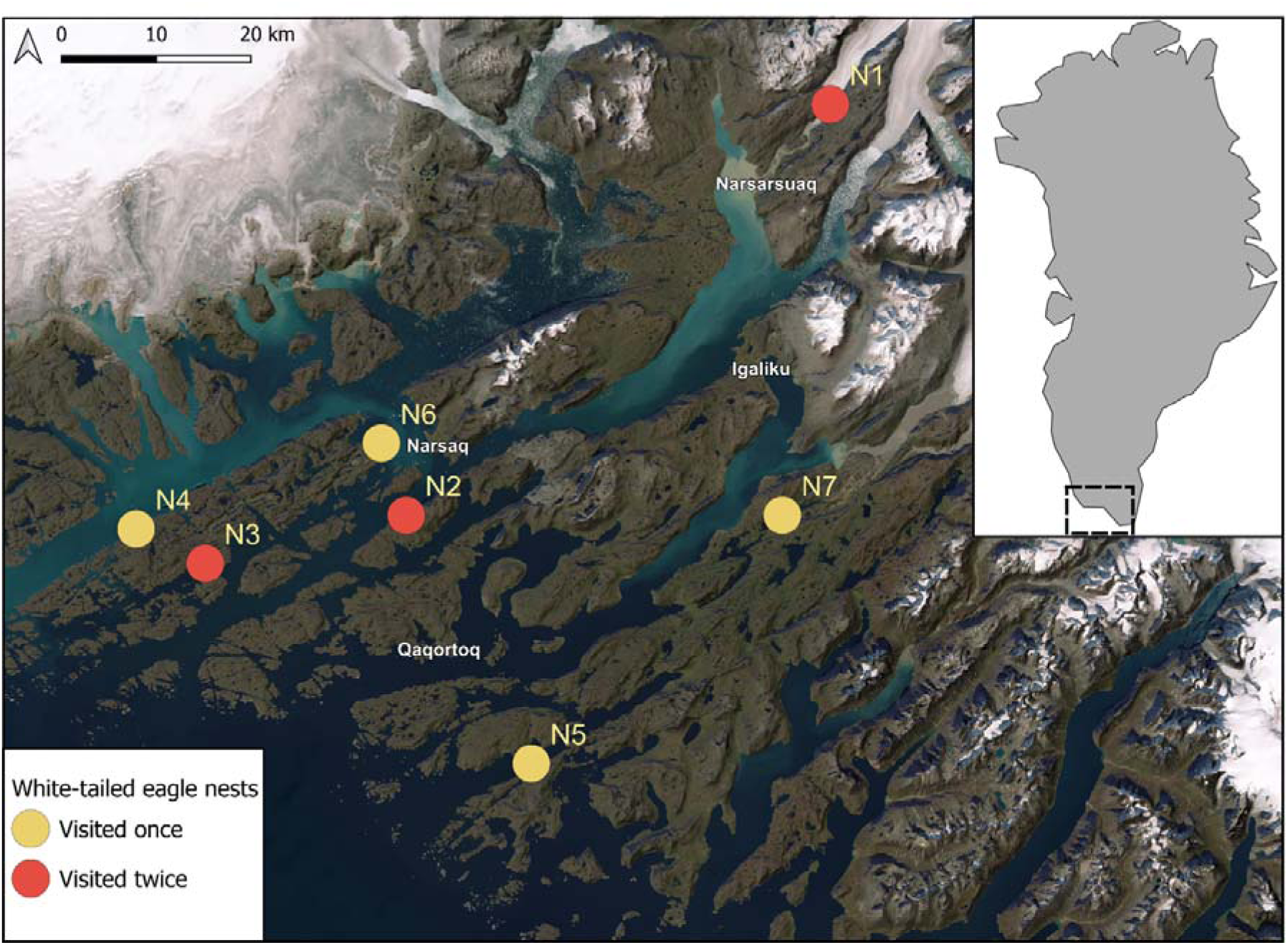
Map depicting the seven white-tailed eagle nests sampled. The inset shows a map of Greenland with the study region outlined by dashed lines. The main settlements of the area are labelled in white. Colours indicate nests: yellow circles: white-tailed eagle nests visited once, red circles: nests visited twice. Points are moved up to 4 km in a random direction to obscure exact locations of breeding sites.

### Sampling

In each nest, we collected faeces, pellets, and soil samples. We also swabbed the beak and all talons of each chick. We collected a total of 99 faeces and 18 pellets from white-tailed eagle nests (Appendix A, Table S1). We sampled faeces using individual tweezers, which we sterilized before use with 5 % bleach. We placed approximately 0.5 mL of faeces into individual 2 mL Eppendorf tubes containing 96 % absolute ethanol. We kept the remaining faecal material in aluminium foil for microplastic assessment. We prioritized sampling fresh faeces based on their availability, focusing on the non-uric (*i.e.,* non-white) portion of the faeces. When a chick defecated during handling, we collected the faecal sample for DNA extraction. We sampled pellets for microplastic assessment, storing them in aluminium foil.

We collected 16 talon and 16 beak swabs from eagle chicks (Appendix A, Table S1). We used nylon bristles moistened in Milli-Q water as swabs, following Bourbour et al. (2019). For beaks (BS), we swabbed the entire external surface of the superior and inferior mandible and the visible skin immediately around the beak. For talons (TS), we swabbed the entire surface of all talons, including toe pads when food remains were visible. We stored the swabs in 2 mL Eppendorf tubes containing 96 % absolute ethanol.

We collected 30 soil samples from eagle nests (Appendix A, Table S1). We collected four soil samples (So) per nest (with six samples in Nest 1), where we only sampled soil during the first nest visit. Each sample consisted of 0.5 ml of soil material placed in individual 2 mL tubes containing 1.2 mL of 96 % absolute ethanol using a teaspoon, which we cleaned and sterilized before use with 5 % bleach. We targeted soil in corners with obvious faecal material, around recent food remains, and in the centre of the nest where the chicks would often be fed.

During fieldwork, we stored DNA samples in the dark at 5 °C, followed by storing them at -25 °C until processing at the Greenland Institute of Natural Resources (GINR, Nuuk, Greenland).

### DNA extraction and PCR

For soil and faecal samples, we centrifuged ∼0.4 ml of material for 5 min at maximum speed and discarded the ethanol supernatant, followed by incubating each sample on a heat block at 56 °C for 40 min to evaporate remnant ethanol. We transferred the dried material into bead beating tubes to extract DNA with the Qiagen DNeasy PowerSoil Pro Kit (Qiagen, Germany), following the manufacturer’s protocol. We included a piece of Mackerel (*Scomber scombrus*) fin as a positive control and PCR grade water as a negative control for each round of extractions. For beak and talon swabs, we similarly removed ethanol but used the Qiagen DNeasy Blood and Tissue kit to extract DNA. Here, we performed an overnight proteinase K digestion at 56 °C. We used agarose gel electrophoresis and the Qubit™ 4 Fluorometer (Thermo Fisher Scientific, Massachusetts, USA) to assess DNA quality and quantity. For swabs, we used a piece of Atlantic cod (*Gadus morhua*) fin as a positive control.

We used a two-step PCR approach to generate metabarcoding libraries. In the first step, we amplified a 100 bp region (V5) of the 12S ribosomal RNA gene using the pan-vertebrate primers described in Riaz et al., (2011) and included Illumina adaptors: Forward - TCGTCGGCAGCGTCAGATGTGTATAAGAGACAGTTAGATACCCCACTATGC, Reverse - GTCTCGTGGGCTCGGAGATGTGTATAAGAGACAGTAGAACAGGCTCCTCTAG. To prevent the amplification of eagle and human DNA, we included a blocking primer for white-tailed eagle (CTATGCCTGGCCCTAAATCTTGATACIIIIICCACCAAAGTATCCGCCTG; Yuan et al., 2024) and *Homo sapiens* (TACCCCACTATGCTTAGCCCTAAACCTCAACAGTTAAATC; Calvignac-Spencer et al., 2013). We divided eagle samples into four different batches. Each batch included a positive control (*i.e.,* a falcon beak swab previously sampled) and a negative control with PCR grade water instead of DNA. We amplified all extracted samples in two replicates that we further pooled.

We amplified DNA as follows: each reaction contained 4 µL of 5x GoTaq flexi buffer (Promega, USA), 1.6 µL of MgCl_2_ (25 mM), 0.4 µL of dNTP mix, 0.1 µL of GoTaq G2 hot start polymerase (Promega, USA), 4.4 µL of PCR grade water, 0.5 µL of pan vertebrate reverse and forward 12S-V5 primers, 2.5 µL of 12S-human blocking primer, 5 µL of white-tailed eagle DPO-blocking primer (Yuan et al., 2024) and 1 µL of extracted DNA, for a total volume of 20 µL. We prepared reactions in sterile PCR tubes (Sarstedt, Germany) and ran PCRs with the following cycles: a 2 min initial denaturation at 94°C, followed by 35 cycles of 10 s of denaturation at 94°C, 30 s of primers annealing at 51°C and 30 s of extension at 72°C, concluded by a final elongation of 10 min at 72°C.

The subsequent indexing PCR, library preparation, and 150 bp paired-end sequencing on an Element AVITI instrument (Element Biosciences, San Diego, USA) was outsourced to the Genomic Technologies Facilities of the University of Lausanne (GTF, Lausanne, Switzerland).

### Bioinformatics and taxonomic assignment

We merged demultiplexed paired-end reads using PEAR 0.9.8 (Zhang et al., 2014) set with a minimum overlap of 20 bp and a minimum PHRED quality score of 25. We trimmed reads using Cutadapt 5.2 (Martin 2011) and removed reads that did not contain both primer sites. We filtered paired reads with VSEARCH 2.28.1 (Rognes et al., 2016) based on a maximum expected error (EE) of 1 and clustered them into Operational Taxonomic Units (OTUs) with more than 97 % sequence similarity (Lucek et al., 2019) and discarded clusters with ≤ 5 reads. We used the uchime3_denovo algorithm implemented in VSEARCH to detect and remove chimeras. Then, we mapped our reads back to OTU centroids with the usearch_global algorithm allowing 97 % sequence similarity. Using a custom Python script, we assigned OTUs to a custom reference database that contained 12S-V5 sequences from the NCBI database (https://blast.ncbi.nlm.nih.gov/Blast.cgi) for all vertebrates occurring in the study area based on literature (Boertmann 1994; Møller et al., 2018; Ugarte et al., 2020; GINR 2026 https://natur.gl/leksikon/land-animals/?lang=en; GBIF https://www.gbif.org/). For taxonomic assignment, we used the blastn tool implemented in BLAST+ 2.16.0 (Camacho et al., 2009) and categorised hits with a query cover < 95 % or an identity percentage < 85 % as unassigned. We assigned hits with an identity percentage ≥ 97 % to the species level, with identity percentage ≥ 93 % to the genus level, and identity percentages ≥ 85 % to the family level (Evans et al., 2025; Wang et al., 2026).

To reduce background sequencing noise, we kept OTUs with a read abundance of > 0.5 % per sample (Paprocki et al., 2024). We were conservative with eagle samples because we detected dietary DNA in some negative controls (see Results). To account for potential lab contamination, we subtracted the read counts of OTUs present in the negative controls from all samples in the corresponding batch, setting any negative value to zero. We compared and curated ambiguous taxonomic assignments using the NCBI database, species occurrence databases, and expert knowledge of local wildlife.

### Microplastic assessment

#### Plastic extraction and detection

We processed white-tailed eagle pellets and faeces in batches of 8, composed of 7 field samples and one blank. First, we kept 1 g of dried samples for plastic extraction. If a sample was not dry, we dried it at 50 °C (Senes et al., 2023). We weighed and processed entire samples that were less than 1 g. We triturated samples in a sterilized fume hood using a glass beaker and then digested them in 40 ml of 10 % KOH at 50 °C for 96 hours at 100 rpm (Provencher et al., 2019; Thiele et al., 2019). Pellets went through a second digestion step with 10 ml 35 % H_2_O_2_ at 50 °C for 24 hours (Charles et al., 2023). Then, all samples went through density separation by adding 200 ml of a 20.5 mol/L NaCl solution to the samples in glass graduated cylinders (Provencher et al., 2019; Thiele et al., 2019). We left samples under the hood to settle for 30 min and filtered the top 40 mL of the decanted solution using vacuum filtration with 11 µm paper filters (Whatman, grade 1, 47 mm) (Dowarah et al., 2020). We prepared a stock solution of Nile red of 1 mg mL^−1^ in acetone and submerged the filters in 10 mL of Milli-Q water. Then, we stained them with 100 μL of the stock solution of Nile red, to obtain a working concentration of 10 μm mL^−1^ (Shruti et al., 2022). We stained filters at 50 °C for 30 min, washed them with Milli-Q water and discarded them. We vacuum filtered each solution with resuspended particles with 0.7 µm glass fibre filters (Whatman, 47 mm). Finally, we dried the filters at 50 °C for 1 h.

We took pictures of the filters using a digital camera (Sony α7 IV, Sony Corp., Tokyo, Japan) with a 35 mm lens. In the dark, we took pictures under a 480 nm forensic blue light (Crime lite 2, FoxFury Lighting Solutions, Oceanside, USA) and placed an orange filter (580 nm) over the lens (Prata et al., 2019). We took a second picture under a 254 nm UV light with the same orange filter (Prata et al., 2019). The camera settings were the following for both wavelengths: F3.5, s1/80, ISO500. Finally, we heated a metallic needle and touched fluorescent particles with it under the 480 nm light. If the particles curled or melted, we confirmed it as plastic (Qaiser et al., 2023).

#### Contamination prevention

To limit and assess the ambient plastic contamination in the lab, we kept all solution in glass containers and covered by aluminium foil. We washed all equipment thoroughly with tap water and rinsed it twice with grade 1 ultrapure water. We performed sample transfers and filtration in a fume hood and cleaned both the hood and the bench between each batch. We wore a white cotton lab coat during the entire process and blue nitrile gloves when required. In addition to procedural blanks done in the lab for each batch, we took a field blank per nest to assess aerial contamination during sampling. It consisted of a 20 x 20 cm sheet of aluminium foil laid on the ground of the nest for the duration of sample collection. We then closed it, and the blank was processed exactly as the other samples for microplastic assessment.

#### Data collection and correction

We analysed each picture twice with the MP-NET software developed by Park and colleagues (2022). First, we annotated all confirmed plastic particles that validated the hot needle test. Confirmed plastic particles accounted for 68.4 % ± 20.4 % of tested particles. When the software did not detect a confirmed plastic particle, we manually added it. Inversely, if the software detected a particle that had not been validated, we manually removed it. The software provided the number of particles and their area. Secondly, we kept all particles detected by the software and, if missing, we manually added the confirmed plastic particles. Those data combined allowed us to have for each sample the number of potential particles, the number of true positive plastic particles (*i.e.,* plastic particles confirmed by needle test), the number of tested particles, the total area of true positive plastic particles and the total area of detected particles. The software failed to detect orange fluorescence of plastic particles under the UV light, because the glass fibre filters emitted unexpected green fluorescence (Prata et al., 2019). Therefore, we excluded from the analysis pictures taken under the UV light.

We performed two corrections. First, 20 % of the filters required a correction because we undertested them with the hot needle compared to the remaining ones. Secondly, we corrected microplastic data for aerial and lab contaminations using data from field blanks. We used Monte Carlo uncertainty propagation (MC) using the R package *tidyverse* (Wickham 2019) to perform both corrections. We used five thousand iterations and the average across blanks for MC correction. Monte Carlo uncertainty propagation MC gives a 95 % probability interval after computing for the correction for the undertested filters, the blanks, and the background noise. Our MC model returned three values: the 2.5 % probability quantile value corresponding to the most conservative value; the 50 % probability quantile value corresponding to the most likely value; the 97.5 % probability quantile value corresponding to the less conservative value.

Using MC corrected data, we first reported the number of samples with a 2.5 % probability quantile value above zero for total plastic area. A MC 2.5 % value above zero means a 95 % probability that the sample has more plastic than the blank and background noise. Then, we reported samples with a MC 50 % values for total plastic area above the limit of detection (LOD) and above the limit of quantification (LOQ) thresholds. LOD is defined as 3x the standard deviation of the blanks for the total plastic area. LOQ is defined as 10x the standard deviation of the blanks for the total plastic area. MC 50 % values are the average values that each iteration of MC will return after correcting for blanks contamination. LOD and LOQ have been recently found to be efficient to correct plastic data with blanks (Waddell et al., 2020; Horton et al., 2021; Dawson et al., 2023). LOD is considered enough to assume presence or absence of plastic contamination (Waddell et al., 2020), whereas LOQ is required for precise quantification of plastic (Horton et al., 2021).

### Statistical analyses

We conducted all analyses in R v. 4.4.3 (R Core Team, 2026) and QGIS 3.34.10 (QGIS Development Team, 2026). We only retained samples with dietary DNA for analysis, excluding four faecal and five soil samples (Table 1a). We calculated the frequency of occurrence (FOO) (O’Hanlon et al., 2025), *i.e.,* the percentage of samples in which a food item was detected for each sample type. We further summarised data at the nest level using the weighted percentage of occurrence (wPOO) and the relative read abundance (RRA) (Yuan et al., 2024; O’Hanlon et al., 2025; Wang et al., 2026). wPOO is FOO rescaled so that the sum of all food items per nest is 100 % and with each occurrence weighted according to the number of taxa in each sample of the same nest. RRA is based on the number of reads of a dietary taxon in a sample, weighted by the total number of reads of the sample. They are calculated following Deagle et al. (2019).

**Table 1:**
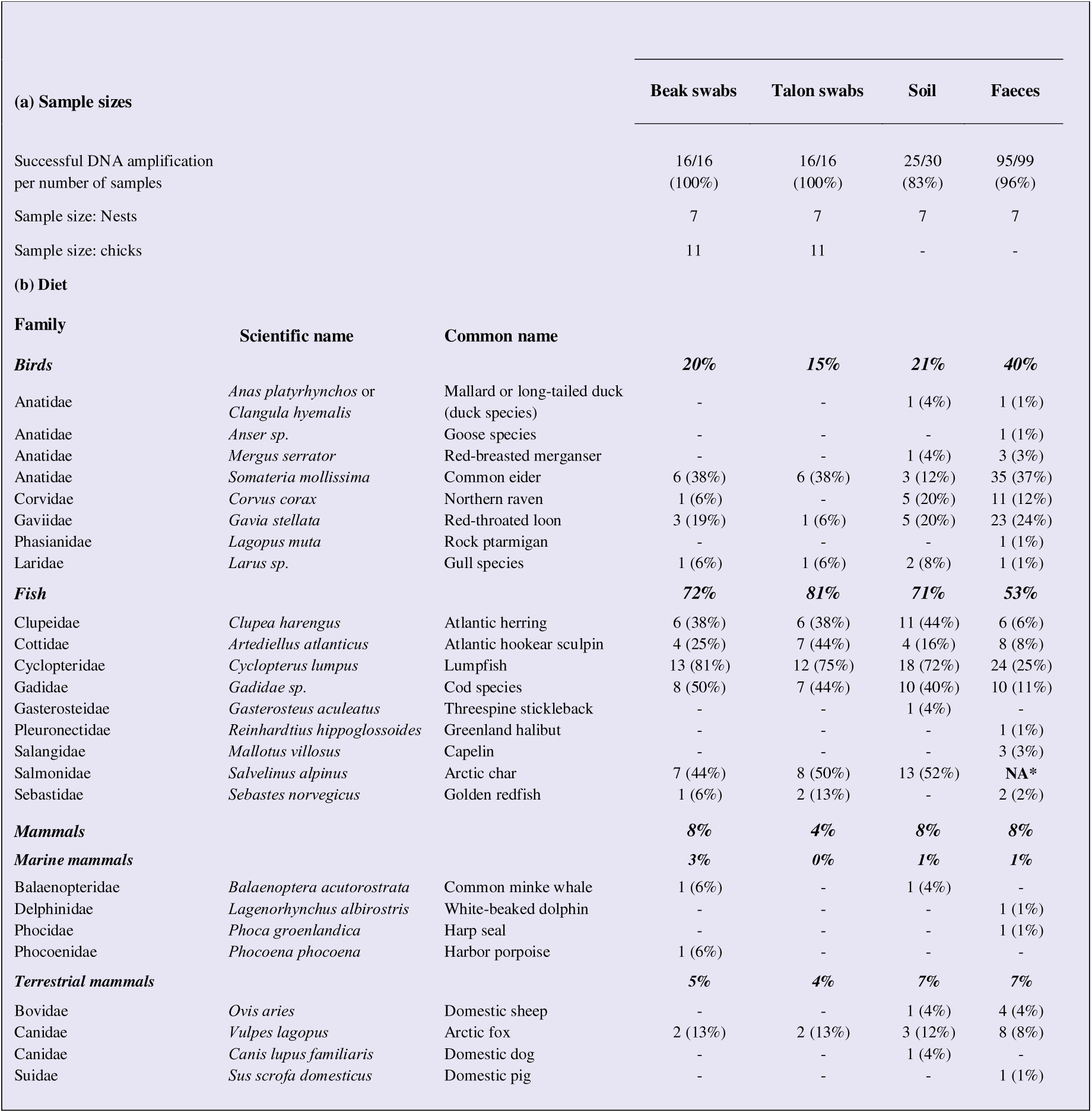
Frequency of occurrence (FOO) of food taxa identified in all sample types of white-tailed eagles (*Haliaeetus albicilla*). Samples were taken in seven different nests in South Greenland during the 2025 breeding period. Only food taxa with > 0.5 % abundance are shown. Arctic char (*Salvelinus alpinus*) was excluded from faeces because of cross-contamination (Appendix A, Fig. S1).

One batch of white-tailed eagle faecal samples was contaminated by a commonly preyed fish, the Arctic char (*Salvelinus alpinus*). The contaminant was too abundant in the negative control to be removed and despite not being present in every sample of the batch (Appendix A, Fig. S1), we decided to exclude this batch from the sample type comparison, hence reducing sample size to 50 faeces from five nests. For the diet analysis, we kept this batch but excluded Arctic char from all faecal samples as it was the only contaminant.

We conducted PERMANOVAs using the *adonis2* function from the R package *vegan* (Oksanen et al., 2026) to test for variation in the diet composition between nest, and sample type (Schwarz et al., 2018; O’Hanlon et al., 2025). We used nest ID and sample type as fixed effects (Höhn et al., 2024). We ran PERMANOVAs based on Jaccard distances for the occurrence data (coded as 1 if present or 0 if absent) and Bray-Curtis dissimilarity for abundance data (RRA; O’Hanlon et al., 2025). We ran the PERMANOVA once including and once excluding faecal samples. We checked the homogeneity of multivariate dispersion assumption for the PERMANOVA with the *betadisper* and *permutest* functions of *vegan*. We visualised community clustering using Non-metric Multidimensional Scaling (NMDS) with the *metaMDS* function implemented in *vegan*. To identify which dietary taxa contributed most to the dissimilarity between sample types, we used a SIMPER similarity percentage analysis, implemented in *vegan* on occurrence data (O’Hanlon et al., 2025).

Using the R package *iNEXT* (Hsieh et al., 2025), we calculated the Hill number across sample types (Wang et al., 2026). By modulating the q parameter of the function, the Hill number either returns species richness (q = 0) or Shannon diversity (q = 1). We explored the relationship between sample sizes of sample types and species richness and Shannon diversity by generating rarefaction and extrapolation curves based on 95 % confidence intervals (Paprocki et al., 2024; Wang et al., 2026).

To explore the difference among sample types in the detection of dietary taxa and to describe eagle diet, we compared the difference in dietary taxa richness obtained in individual samples per sample type using a GLMM, implemented in the R package *glmmTMB* (Brooks et al., 2017). The model used dietary richness per sample as response variable and sample type (TS, BS, FA, So) as a fixed effect with nest ID as a random intercept, implementing a truncated negative binomial distribution. We used a likelihood ratio test where the GLMM was tested against a null, or intercept only, model (Tredennick et al., 2021). We tested GLMM assumptions with the R package *DHARMa* (Hartig et al., 2024). We used Tukey tests from the R package *emmeans* (Lenth et al., 2026) to determine pairwise differences among sample types.

To identify whether one or more dietary taxon was associated with plastic pollution in faecal samples, we performed an Indicator Species Analysis and species by species Fisher’s Exact Tests using the R function *fisher.test* (Höhn et al., 2024), where species composition was the response and plastic contamination the explanatory variable. We performed those tests on occurrence data from all faecal samples that were screened for microplastic and that contained dietary DNA after excluding Arctic char from the analysis. We defined plastic contamination categorically: samples contaminated with plastics returned TRUE (*i.e.,* area MC 50 % values > LOD), while those without plastic returned FALSE (*i.e.,* area MC 50 % values < LOD).

## Results

We detected dietary DNA in 25 out of 30 (83 %) soil samples, in 95 out of 99 (96 %) faeces, and in every other sample (Table 1a). However, the detection rate in faecal samples may be overestimated because five of them only contained Arctic char that could also be attributed to contamination. Soil samples generally yielded fewer read numbers compared to other sample types.

### White-tailed eagle diet

Using metabarcoding data, we detected 25 dietary taxa across the four sample types (Fig. 2, Table 1b). Of those, nine were fish, eight birds, and eight mammals (terrestrial and marine). The primers used did not allow to differentiate closely related species for several genera including the bird genuses *Larus, Anser,* and the fish family *Gadidae*. The most common dietary taxa by read abundance (Fig. 3) and occurrence data (Table 1b; Appendix A, Fig. S2) were lumpfish (*Cyclopterus lumpus*), cod species (*Gadidae*), and Arctic char. Atlantic herring (*Clupea harengus*) was more common in the occurrence data (Table 1b; Appendix A, Fig. S2). Fish are the most common group of dietary taxa. Birds are the second most identified group (Fig. 3, Table 1b; Appendix A, Fig. S2), especially the common eider (*Somateria mollissima*). In faeces, other important bird species are red-throated loon (*Gavia stellata*) and northern raven (*Corvus corax*) (Fig. 3A; Appendix A, Fig. S2A). Mammals were the least common group (Fig. 3, Table 1b; Appendix A, Fig. S2), with Arctic fox (*Vulpes lagopus*) being the most common mammal found across all samples. Based on soil samples and beak and talon swabs (Table 1b), fish accounted for around 75 % of all dietary occurrences, birds for 19 %, and mammals for 7 % of all dietary occurrences (Table 1b; fish: So = 71 %, BS = 72 %, TS = 81 %; birds: So = 21 %, BS = 20 %, TS = 15 %; mammals: So = 8 %, BS = 8 %, TS = 4 %). For faeces, we had to apply the average occurrence of Arctic char from other sample types (mean FOO = 49 %; Table 1b) to calculate the occurrence of each dietary group. This was due to the contamination from Arctic char in some faeces. Fish then accounted for 53 % of all dietary occurrences in faeces, birds for 40 %, and mammals for 8 %.

**Figure 2:**
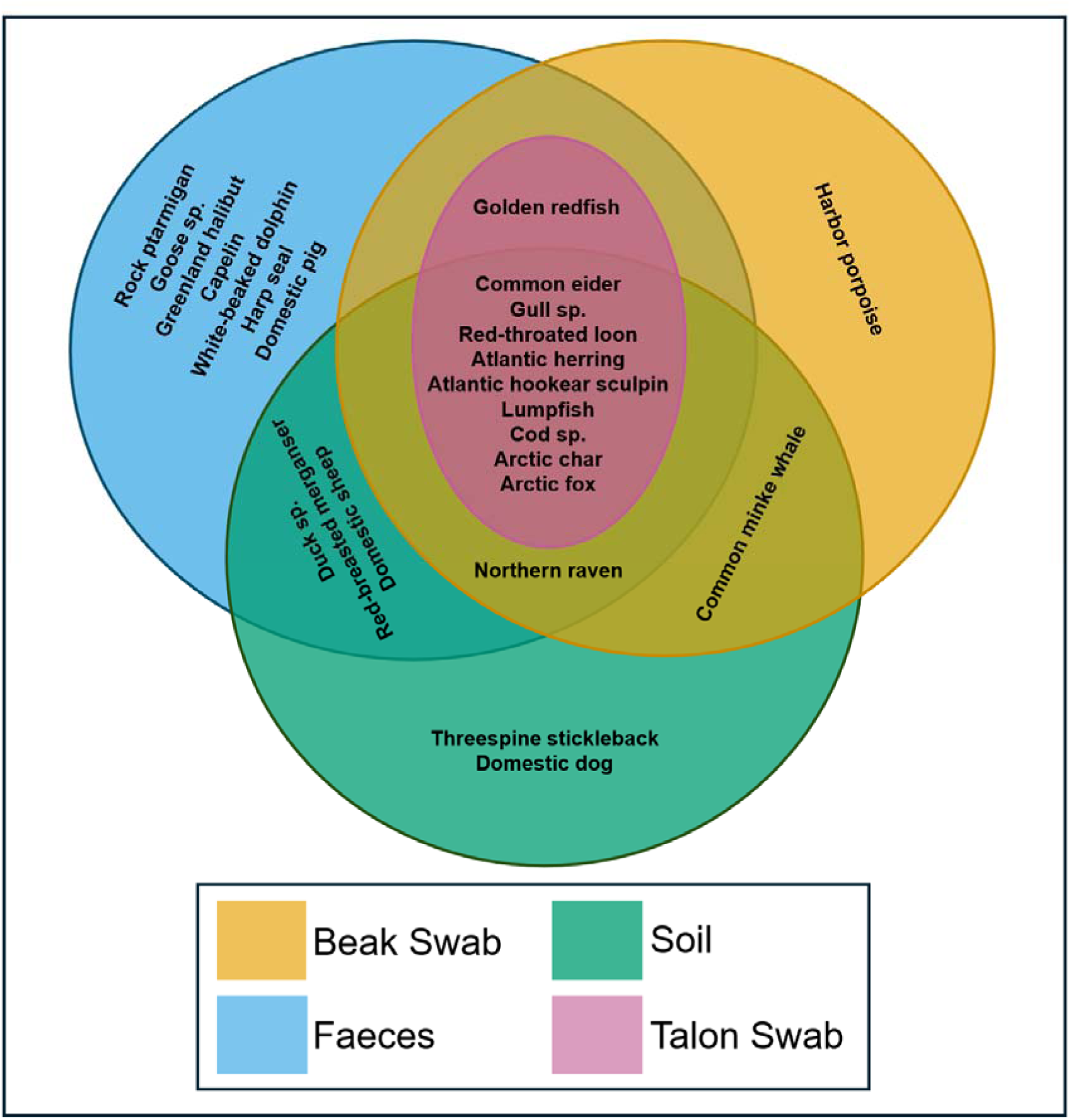
Venn diagram of white-tailed eagle (*Haliaeetus albicilla*) dietary taxa identified using each sample type. Colours indicate sample type: orange– beak swabs (BS), blue – faeces (FA), green– soil samples (So) and purple – talon swabs (TS).

**Figure 3:**
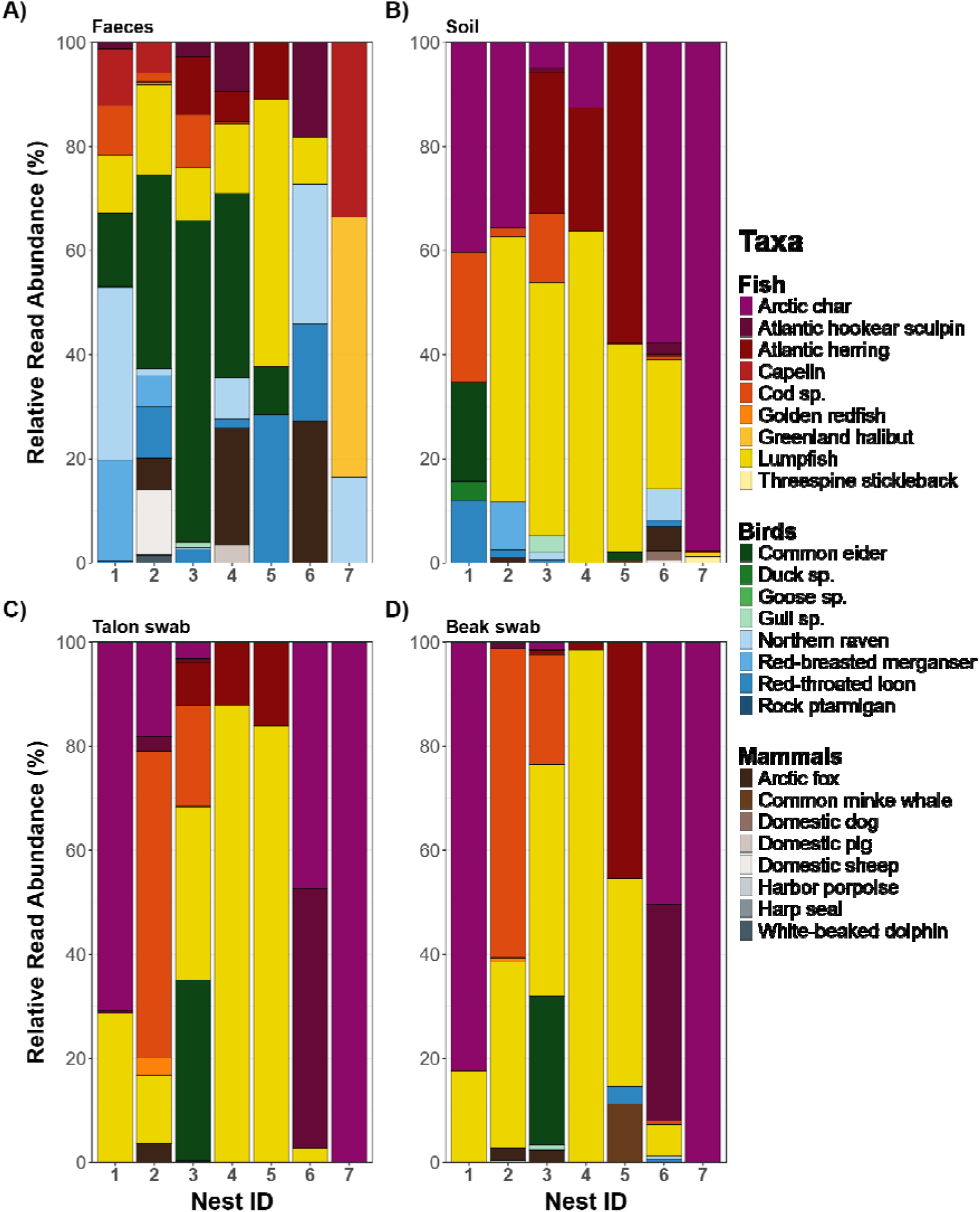
Relative read abundance (RRA) in percent for each dietary taxon from white-tailed eagle sample types at each nest visited during the 2025 breeding season. A) Dietary taxa in faeces, excluded Arctic char due to cross contamination. B) Dietary taxa in soil samples, C) talon swabs and D) beak swabs.

### Effect of sample type on diet description

Extrapolation curves for Hill numbers based on eagle talon swabs reached a plateau indicating that the sample size is sufficient to capture the diet (Fig. 4). Other sample types approached saturation for the Shannon index (q=1) (Fig. 4B). However, beak, soil, and faecal samples indices continued to show an increasing trend which is emphasised by dietary taxa richness (q=0) (Fig. 4A), suggesting that additional sampling would provide additional species richness and, to a lesser extent, species diversity in the eagle diet. Beak swabs and nest soil captured more dietary taxa than faeces and talon swabs for the same sample size (Fig. 4A-B).

**Figure 4:**
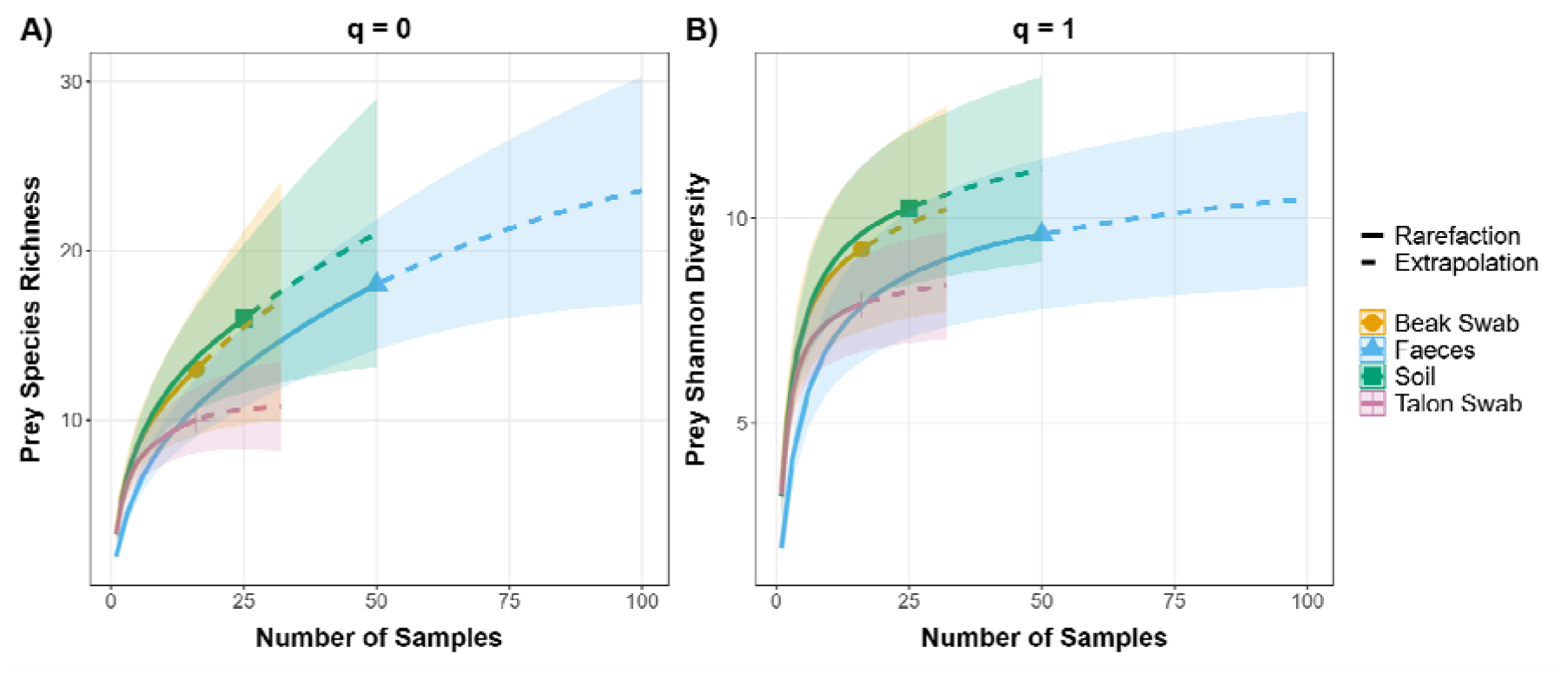
Hill number diversity rarefaction curves (solid lines) for white-tailed eagle and peregrine falcon diet with their extrapolation (dashed lines) based on 95 % confidence intervals. A) Dietary taxa richness rarefaction and extrapolation curves (q = 0) of the different white-tailed eagle sample types. B) Dietary Shannon diversity rarefaction and extrapolation curves (q = 1) of the different white-tailed eagle sample types. Only faecal samples that were not contaminated with Arctic char were used (N = 50).

Dietary composition varied significantly between nests and sample types for both RRA (PERMANOVA; Nest: *F*_6, 96_ = 8.87, *R*^2^ = 0.34, *p* = 0.001; Sample type: *F*_3, 96_ = 1.96, *R*^2^ = 0.04, *p* = 0.025; Appendix A, Fig. S3) and species occurrence (Nest: *F*_6, 96_ = 9.84, *R*^2^ = 0.36, *p* = 0.001; Sample type: *F*_3, 96_ = 2.70, *R*^2^ = 0.05, *p* = 0.002; Appendix A, Fig. S3). When excluding faecal samples, dietary composition varied significantly between nests for both RRA (Nest: *F*_6, 48_ = 7.73, *R^2^* = 0.49, *p* = 0.001) and species occurrence (Nest: *F*_6, 48_ = 9.44, *R*^2^ = 0.53, *p* = 0.001), but sample types did not have a significant effect on dietary composition for both RRA (Sample type: *F*_3, 48_ = 0.49, *R^2^* = 0.01, *p* = 0.848) and occurrence (Sample type: *F*_3, 48_ = 1.08, *R^2^* = 0.02, *p* = 0.359). Nest explained consistently more variation in diet than sample type based on *R^2^* values. Moreover, only faeces brought significant variations of diet between sample types. Multiple dietary taxa explained the difference of diet between sample types (SIMPER test; all *p* < 0.05; Table 2).

**Table 2:**
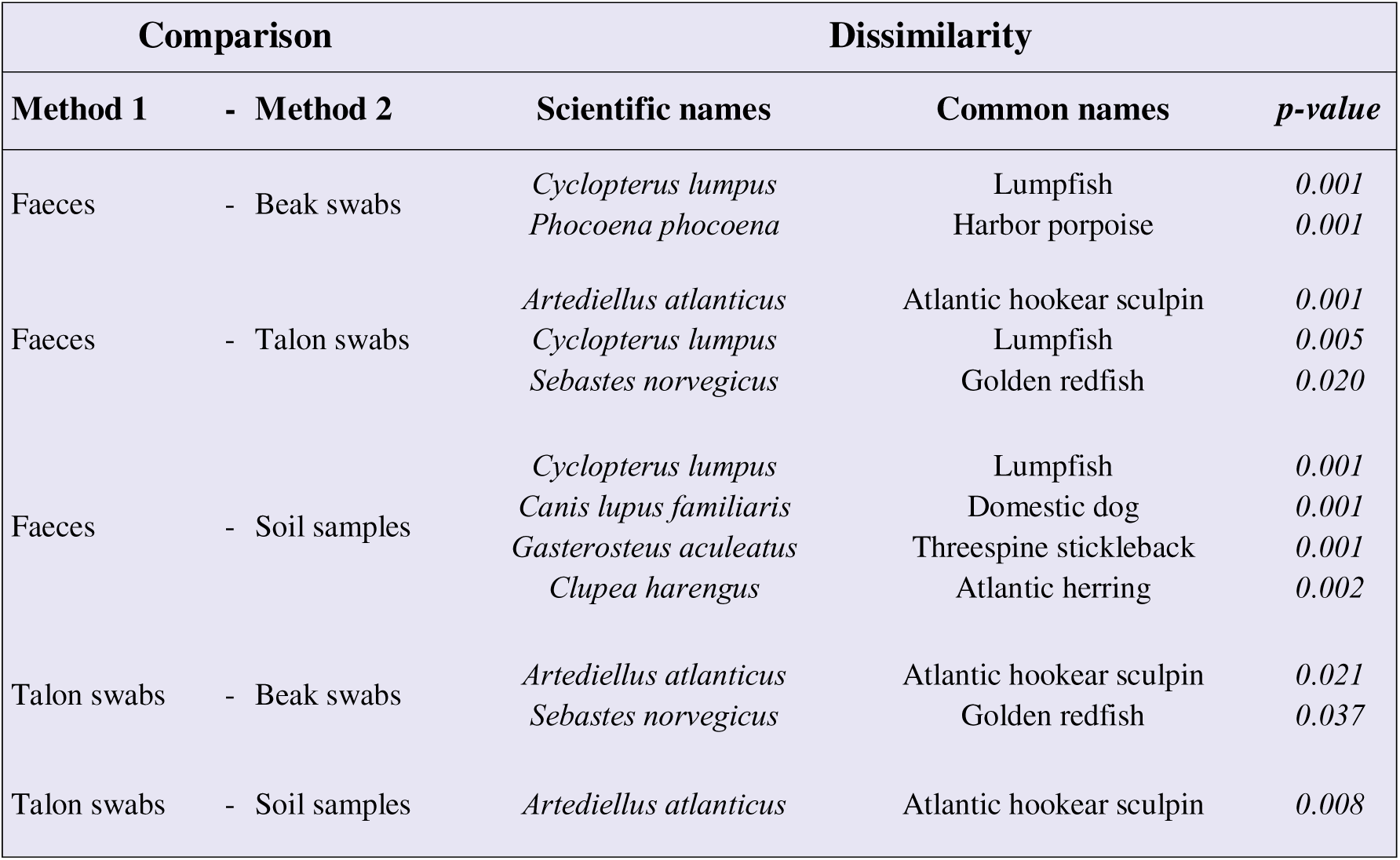
Dietary species driving significant dissimilarities between white-tailed eagle sample types based on SIMPER tests on occurrence data that excluded contaminated faecal samples.

### Dietary detection rates among sample types

The number of dietary taxa detected per sample differed significantly among sample types (GLMM, Likelihood Ratio Test: = 24.64, *p* < 0.001; Fig. 5). Faeces detected significantly fewer taxa per sample than beak swabs (Tukey post-hoc pairwise comparison: *p* = 0.002), talon swabs (*p* = 0.004), and soil samples (*p* < 0.001; Fig. 5). Dietary richness did not statistically differ between beak swabs, talon swabs, and soil (all *p* > 0.05; Fig. 5).

**Figure 5:**
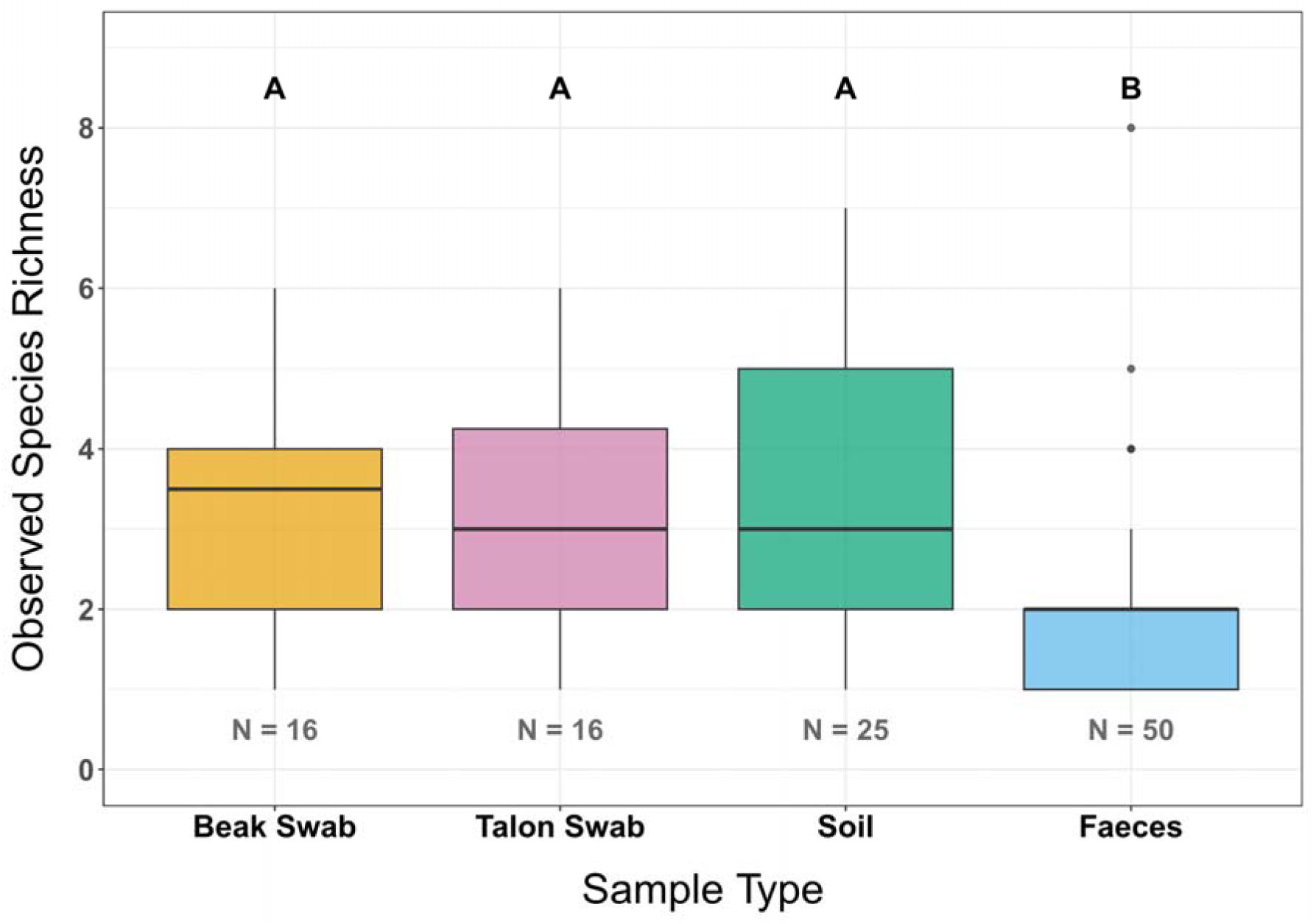
Observed dietary taxa richness of samples that contained dietary DNA per sample type. Distribution of dietary richness found in each sample type. Types without a letter in common significantly differ statistically. Sample sizes are indicated for each type.

### Microplastic association with diet

We analysed 76 faeces and 21 pellets from six nests (access to the Nest 6 caused logistical limitations, hence preventing data collection) and found microplastics in all samples and blanks. Of those 93 samples, 21 faeces) and ten pellets, for a total of 31 samples (33 %), were above the 95 % probability threshold set by the Monte Carlo uncertainty propagation with total plastic area per sample (MC 2.5 % values > 0; Appendix A, Fig. S4). Only Nest 7 showed no statistical evidence for microplastics. 14 faeces and 7 pellets, for a total of 21 samples out of 93 (15 %) were above the limit of detection LOD (MC 50 % values > LOD; Appendix A, Fig. S4). A total of four samples, all faeces, out of 93 (4 %) were above the limit of quantification LOQ (MC 50 % values > LOQ; Appendix A, Fig. S4). After removing Arctic char from analysis, no dietary taxa were significantly associated with microplastic contamination (Fisher’s exact test; all p > 0.05; Appendix A, Fig. S5).

## Discussion

Classic methods to study diet are often time consuming and rely on human skills for dietary identification (Nielsen et al., 2018), while some dietary taxa are impossible to detect visually (*i.e.*, chunks of meat) (Wille and Kampp 1983). We visited seven nests of white-tailed eagles in 17 days, allowing us to sample 11 eagle chicks in South Greenland. Using a metabarcoding approach, we successfully described the chicks’ diet. We compared the efficacy of different sample types. With a total of 25 dietary taxa, the eagle diet was mainly composed of fish (Table 1). We uncovered a broader diet for our focal species than previously reported from Greenland (Wille 1979; Wille and Kampp 1983). Our four sample types detected important dietary information, although faeces provided a different overview of the diet than the other sample types (Table 1b; Appendix A, Fig. S3). We discuss these findings in the broader context of the biology of our focal species and the utility of different sample types coupled with metabarcoding to study diet.

### Feeding ecology of the white-tailed eagle

Former studies on the white-tailed eagle showed contrasting results, where fish were sometimes the least and birds the most common dietary group (Whitfield et al., 2013; Reid et al., 2023; Yuan et al., 2024). As the white-tailed eagle is an opportunistic and generalist bird of prey (Nadjafzadeh et al., 2016a), differences in dietary composition can be explained by regional variation in prey abundance and availability, where fish are more present in the diet of individuals near water bodies compared to inland areas (Reid et al., 2023). Consistently, we found the diet to differ between nest sites based on available resources, where Arctic char was the dominant food item in nests near freshwater bodies and common eider was more important in nests near the outer coast (Appendix A, Fig. S6). Overall, fish were the main dietary group of the white-tailed eagle, followed by birds and mammals (Table 1b), which was similar to the diet preference of white-tailed eagles in Sweden (Ekblad et al., 2020). This is consistent with earlier studies on chick diets of this species in Greenland using camera recordings and direct observations, which showed that fish dominated the diet up to 90 %, while birds and mammals accounted for the remainder (Kampp and Wille 1979; Wille 1979; Wille and Kampp 1983).

While the V5 region of the mitochondrial 12S RNA (Riaz et al., 2011; Yuan et al., 2024; Wang et al., 2026) is appropriate for diet analysis of digested and degraded template material, we observed a limited taxonomic resolution for closely related species such as birds of the *Larus* genus and *Anatidae* family, and fish from the *Gadidae* family. For these groups, we obtained the same identity and coverage scores to the species level for more than one likely prey species. Specifically, mallard (*Anas platyrhynchos*) reads were equally assigned to long-tailed duck (*Clangula hyemalis*) but not to any other species of the *Anatidae* family. Both mallard and long-tailed duck are present in the study area and are likely prey species of the white-tailed eagle. We therefore discuss this taxonomic conflict as duck sp. This and the other taxonomic conflicts could lead to underestimating species richness and diversity of the studied diet.

The three-spine stickleback (*Gasterosteus aculeatus*) and the Atlantic hookear sculpin (*Artediellus atlanticus*) are small fish, the sculpin being also benthic. Those species were likely picked up through cods or Arctic char through secondary predation (de Bruyn et al., 2021; Good et al., 2024; O’Hanlon et al., 2025). Similarly, while capelin (*Mallotus villosus*) and Atlantic herring (*Clupea harengus*) are prey of cod and Arctic char to some extent, eagles will also actively feed on these two smaller fish species (Wille and Kampp 1983; Kampp and Wille 1997).

In Greenland, the only mammal consistently present in white-tailed eagle chick diets was the Arctic fox (Wille and Kampp 1983; Kampp and Wille 1997). We detected seven more mammal species in addition to the Arctic fox, which aligns with identification of food items from nests that suggest that eagles may consume other mammals in Greenland, such as Arctic hares, domestic sheep or lambs, and reindeers (Kampp and Wille 1979), while historical texts suggest the list could be expanded further with small seals (Egede 1770). We found presence of large marine mammals in the diet, which was rare and could only be explained by either scavenging or secondary predation by the Arctic fox, raven or gulls (de Bruyn et al., 2021; Good et al., 2024; O’Hanlon et al., 2025). Eagles could also have fed on carrion or waste originating from fishing and hunting activities, as local inhabitants are allowed to hunt whales and other marine mammals. This suggests that eagles do not only feed on carrion during the winter season as hypothesised by Nadjafzadeh et al., (2016b), but also during the breeding season. Similarly, domestic pigs (*Sus scrofa domesticus*) are not raised in Greenland, and eagles may have fed on waste from human consumption, or it may be secondary predation picked up through another scavenger like the Arctic fox. The occurrence of domestic sheep (*Ovis aries*) in the white-tailed eagle diet was very low (Table 1b). Here, domestic sheep DNA in soil samples was likely due to the use of wool as nest material. Sheep DNA in faeces unlikely comes from secondary predation as Arctic fox and northern raven were absent from samples containing sheep. Thus, in faeces, sheep DNA likely derived from ingested tissues (Madsen and Vibe 1950; Kampp and Will 1979). We are unable to identify whether the tissue originated from carrion, or depredation, but ingestion of nest material (sheep wool) seems unlikely to explain the obtained sheep DNA in faeces as raptors do not digest keratin and regurgitate hair in pellets (Leprince et al., 1980; Duke 1997; Houston et al., 2007). Domestic dog (*Canis lupus familiaris*) was another rare species whose presence is possibly due to predation, feeding on carrion, or secondary consumption. Dogs occur in the study area as working animals at farms and as pets.

In general, DNA metabarcoding cannot identify whether the DNA sampled comes from actively hunted prey, consumption of carcasses, or whether the DNA comes from another source, such as through secondary predation or generally from the environment, *i.e.,* nest material. Additionally, as we only collected samples from chicks during the breeding season, we cannot generalize on the eagle’s diets beyond this life stage.

### Dietary information among sample types

Overall, we succeeded in recovering DNA from all four sample types (Table 1). The different sample types provided complementary dietary information (Fig. 2), although with different efficacy. Importantly, all DNA sample types are sensitive to cross and exogenous contaminations (Appendix A, Fig. S1) (Deagle et al., 2019), *e.g.,* picking up non-dietary DNA through contact with the substrate. Additionally, PCR bias in sequence amplification and differences in DNA quantity per dietary taxa can also lead to recovery biases (Deagle et al., 2019). Now, talon swabs captured diet adequately with the available sample size (Fig. 4), whereas the other sample types did not reach saturation, suggesting that additional samples would provide further dietary information.

We failed to recover dietary DNA in some faeces (Table 1a), likely because faeces are composed of body fluid containing a substantial amount of host DNA (Paprocki et al., 2024). Our detection rates for faeces were comparable to other studies (98 %, Trevelline et al., 2018; 95 %, Nørgaard et al., 2021; 94 %, Wang et al., 2026). We also failed to recover dietary DNA in some soil samples where read numbers varied greatly (Appendix A, Table S2), likely because of the heterogeneity of the nest material and the limited potential of our extraction protocol on some food remains, such as feathers. In feathers, DNA is encapsulated in a keratin shell (Horváth et al., 2005) that could limit the transfer of dietary DNA to the soil and may thus not have been recovered with the soil extraction protocol used. Therefore, we propose that a food item type and/or its size could be associated with differences in read numbers and hence its detectability in soil samples (Deagle et al., 2019).

Beak and talon swabs had the best detection rate across sample types, consistent with a former study on the rough-legged hawk (*Buteo lagopus*) (Paprocki et al., 2024). Beaks and talons of white-tailed eagles generally showed little to no food residue, possibly because of post-eating cleaning behaviours (pers. obs. from authors). Nevertheless, our results demonstrate that swabbing external surfaces is also suited to larger raptors than previously studied (Bourbour et al., 2019; Bourbour et al., 2021; Paprocki et al., 2024). A previous study showed comparable efficiency at identifying dietary richness and diversity between swabs, with talon swabs detecting more dietary taxa than beak swabs (Paprocki et al., 2024). Conversely, we detected overall slightly less dietary taxa for talon compared to beak swabs (Fig. 4A-B), with talons missing out some rare food taxa. Other studies that collected swabs focused on fledged raptors during migration, suggesting talons to provide more dietary information for raptors old enough to hunt their prey than for young chicks (Bourbour et al., 2019; Bourbour et al., 2021; Paprocki et al., 2024). Focusing on raptor chicks, talon swabs were likely less adequate than beak swabs. As chicks are likely to be assisted by parents when feeding on large and/or avian food items, they do not use their talons to feed on all items (Kampp and Wille 1997; Sonerud et al., 2014). The beak being the primary contact point with food could explain why uncommon avian and mammalian taxa are more present in beak than talon swabs in comparison to fish taxa (*i.e.,* soft and easy-to-eat tissue). Despite their efficacy, both talon and beak swabs require handling the birds. It is also challenging to collect swabs in large sample sizes as large raptors such as eagles are scarcely distributed, have large territories, and have small brood sizes.

Faecal samples captured more dietary taxa overall than any other sample type; this likely reflects the larger sample size, because at matching number of samples, faeces obtained a lower dietary diversity and number of taxa (Fig. 4A-B). Overall, we detected fewer dietary taxa per faecal sample than with other sample types (Fig. 5). Faeces typically reflect food consumed shortly before defecation (Schattanek et al., 2021), but traces of DNA can be found up to a few days after ingestion (Deagle et al., 2010). Birds’ high frequency of defecation coupled with the low richness recovered by individual faeces supports the observation of Paprocki et al., (2024) that faeces contain dietary DNA from shorter pre-sampling intervals than swabs. Faecal samples recovered fewer fish taxa in both FOO (Table 1b) and RRA (Fig. 3), but more bird taxa, which is similar to a metabarcoding study on white-tailed eagle faeces in China (Yuan et al., 2024). This difference was not explained by the exclusion of Arctic char, as the main fish taxa detected by other sample types were equally less prevalent in faeces (Table 1b). Soft tissues, as from fish, may be more efficiently digested than other food tissues, passing more quickly through the digestive system and could therefore account for the reduced detectability (Thomas et al., 2014; Schattanek et al., 2021; Paprocki et al., 2024). Therefore, faeces require larger sample sizes, may underestimate the importance of fish in diets, and mainly detect recent meals, despite being non-invasive, and opportunistically collected samples (O’Hanlon et al., 2025).

Soil from eagle nests captured the highest number of dietary taxa among all sample types at matching sample sizes, but likely not all food items were recovered (Fig. 4A). Soil can show dietary information from food ingested over a longer period than faeces (Good et al., 2024). Our results are consistent, as dietary richness per sample was higher for soil than faeces (Fig. 5), giving comparable dietary information to beak and talon swabs. For example, we found the presence of the common minke whale (*Balaenoptera acutorostrata*), a rare dietary species, in a beak swab and soil originating from the same nest. However, soil samples may contain DNA originating from nesting material, as illustrated by sheep DNA detected in an eagle nest containing sheep wool. Despite that, sampling soil material from nests appears a promising, yet not often implemented approach to study raptor diets.

### Microplastics and diet

We found that microplastic contamination in our study area was present but low (Appendix A, Fig. S4), often being close to our significance thresholds. Information on the presence of microplastics in trophic interactions in Greenlandic ecosystems is rare, but studies reported contamination of two seabirds: the little auk and the thick-billed murre (Provencher et al., 2014; Amélineau et al., 2016). White-tailed eagles are at the top of the trophic chain in South Greenland, connecting both marine and terrestrial ecosystems. A low plastic contamination could indicate low contamination across lower trophic levels (Deoniziak et al., 2022; Nessi et al., 2022). However, microplastics larger than 200 µm can decrease along the food chain, whereas plastics smaller than 100 µm are likely to biomagnify (Covernton et al., 2022; Qaiser et al., 2023). Against our prediction, none of the consumed fish species was associated with plastic contamination. Variations in processing time of microplastics and food items by the digestive system could create a temporal mismatch between the detection of microplastics and dietary DNA in faeces, but no data to our knowledge currently exist to support this hypothesis.

## Conclusion

Taken together, we demonstrate the value of samples from faeces, nest soil, beaks, and talons in identifying diet using DNA metabarcoding. Our results show that eagles fed their chicks mainly fish, fewer birds, and even fewer mammals. We observed substantial variation in dietary taxa among eagle nests, demonstrating the influence of local food availability on diet. Faeces were the only sample type that provided significantly different dietary information with a reduced presence of fish (Fig. 3, Table 1b; Appendix A, Fig. S2). Our results further show that all sample types successfully recovered dietary DNA, with soil samples being most promising for eagles. Soil and beak swabs gave comparable results, while talon swabs missed some rare avian and mammalian items, and faeces required an increased sample size. Finally, we reveal that the contamination of white-tailed eagles by microplastics is low but present in South Greenland, even though no food species could be identified as a sole vector of plastic pollution. Further research is needed to identify the duration for which DNA from different food taxa can be detected in nest soil. Determining how digestion affects detection of different tissue consistencies and dietary groups in faecal samples is also needed. Our finding of microplastics, highlights the need for additional data to assess their impact and accumulation across the Arctic food chain.

## Acknowledgments

We thank BlueIce, Riding Greenland, and residents of Narsarsuaq and Qanisartuut for logistical support. All fieldwork occurred with permission from the Government of Greenland (permits: 25328513, G25-029). This project was supported by the Greenland Research Council (to NPH J.nr. cd339), the Aage V. Jensen Fonde (to NPH), the Swiss Polar Institute (nr. INT-2025-01 to MP), the Basler Stiftung für biologische Forschung (to MP), the Fonds des donations of the University of Neuchâtel, the Société Zoologique de Genève (to MP), and the Sophie Danforth Conservation Biology Fund (SDCBF to MP). KL was supported by the Swiss National Science Foundation grant 202869.

## Data availability statement

Sequence data is deposited on NCBI (Bioproject PRJNA1533926, https://www.ncbi.nlm.nih.gov/sra/PRJNA1533926). All scripts will be deposited on Zenodo following acceptance.

## Author contributions

MP, KL, and NPH conceived and designed the study. MP, NPH, and KF collected the data. MP analysed the data. All authors interpreted the results. MP wrote the first draft. All authors contributed to the final draft.

## Permission statement

No material from other sources has been reproduced.

## Appendix A

### Supplementary figures

**Figure S1:**
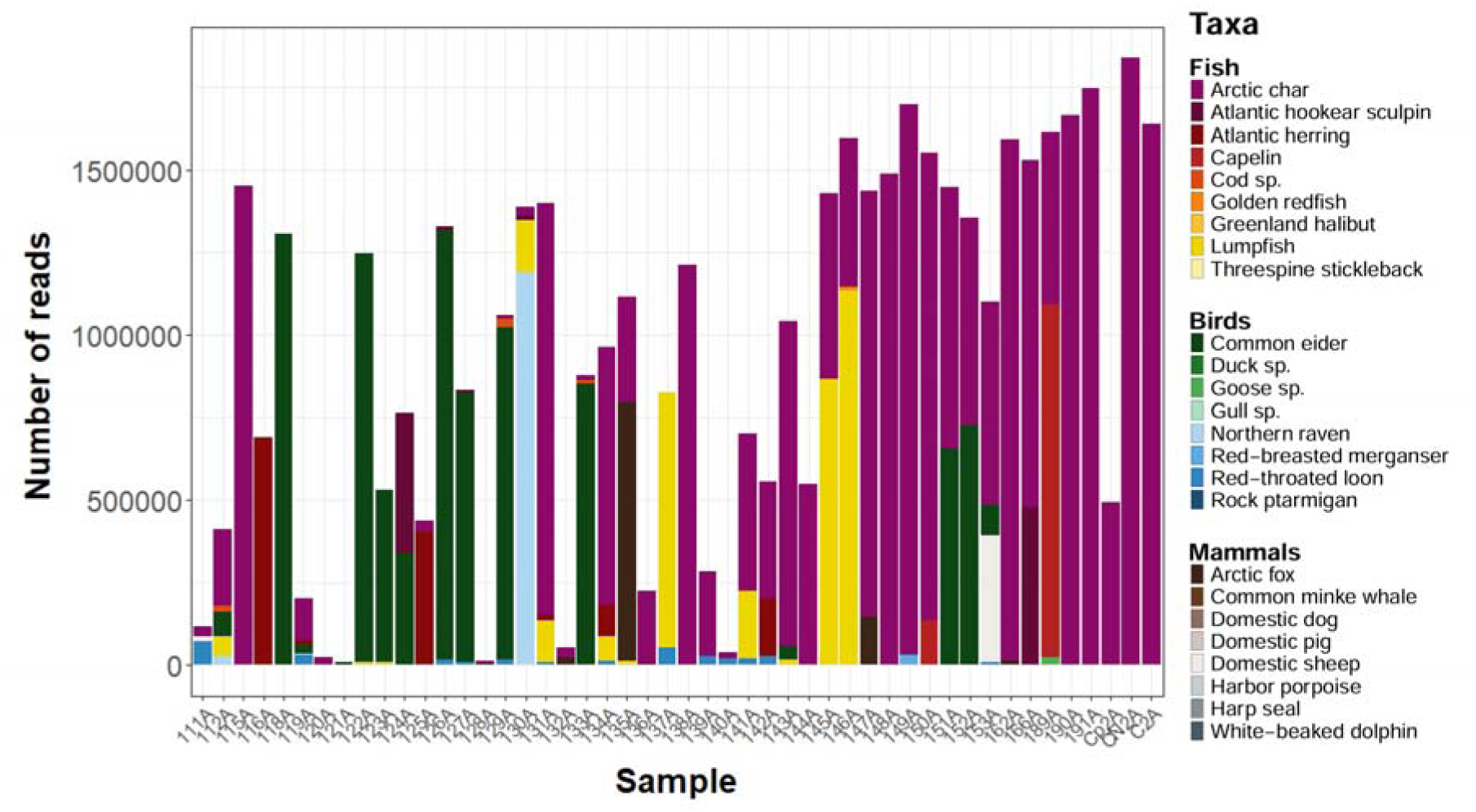
Number of reads present in every sample of the white-tailed eagle PCR batch 2. This batch contained only faecal samples and showed cross-contamination of Arctic char, *Salvelinus alpinus* in yellow. CN2A and C2A are the negative controls of this batch. Cp2A is the positive control that contained DNA from a peregrine falcon’s beak swab previously analysed. The positive control also displayed contamination by Arctic char that should be absent from this sample.

**Figure S2:**
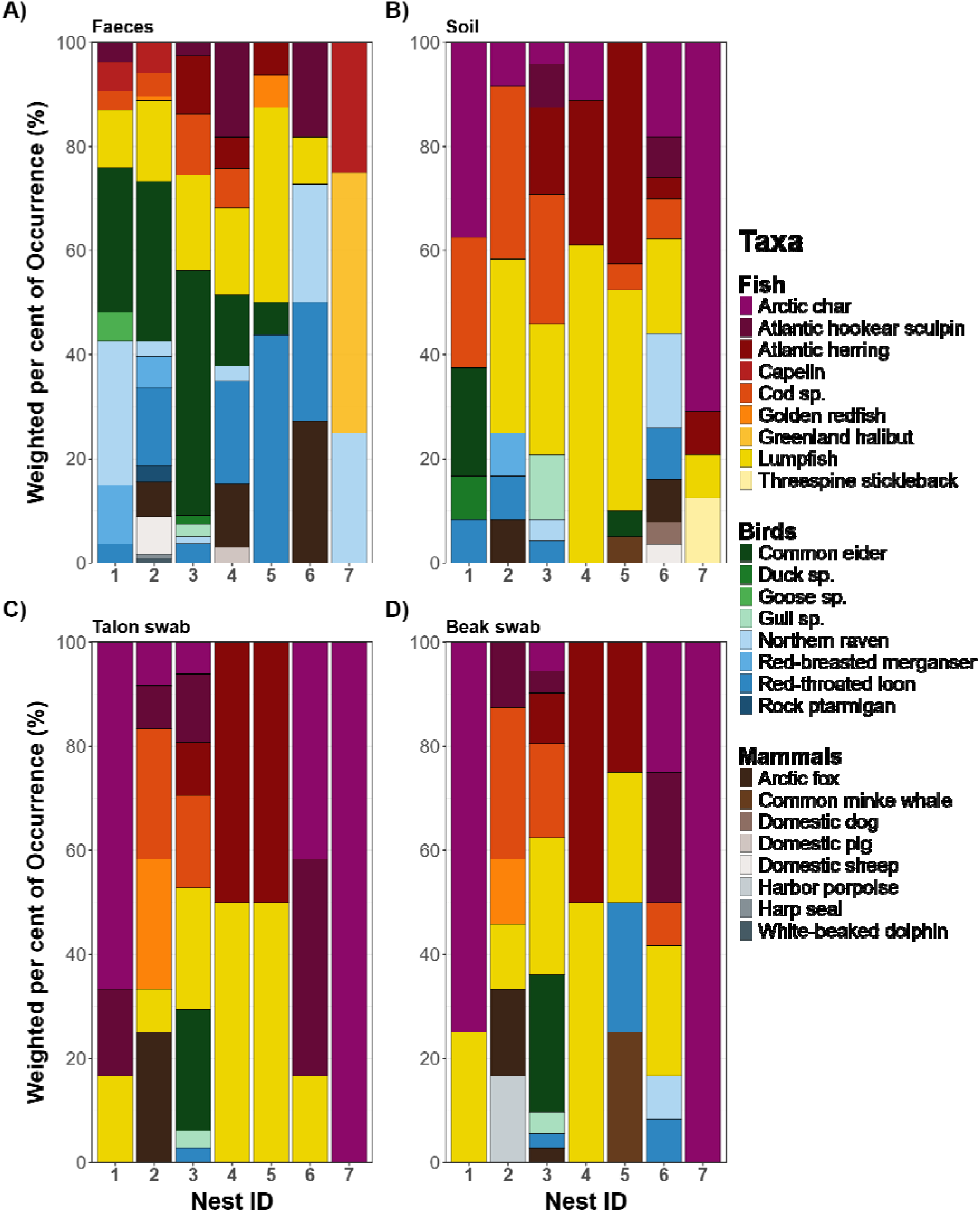
Weighted percentage of occurrence (wPOO) for each dietary taxon from white-tailed eagle sample types at each nest visited during the 2025 breeding season. A) Dietary taxa for faeces, excluded Arctic char due to cross-contamination. B) Dietary taxa in soil samples, C) talon swabs and D) beak swabs.

**Figure S3:**
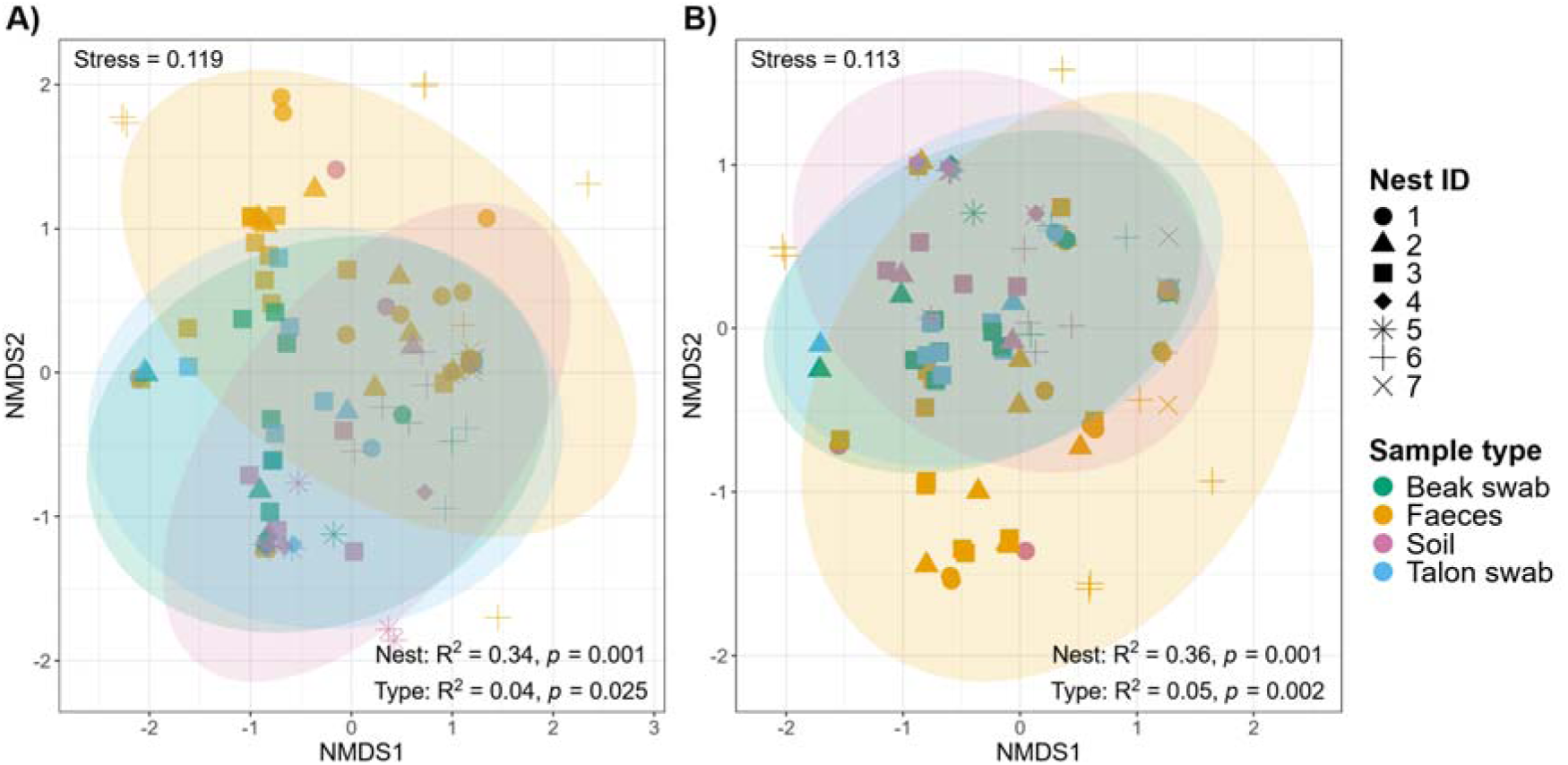
Comparison of sample types, including uncontaminated faeces, between white-tailed eagle nest ID using a Non-metric Multidimensional Scaling (NMDS). Left, NMDS based on Bray-Curtis dissimilarity on relative read abundance (RRA) of food items in each sample, stress-value = 0.12. Right, NMDS based on Jaccard distance based on presence/absence of food items in each sample, stress-value = 0.11. Each nest ID is differentiated by shape. Colours indicate sample type: red – faeces (FA), green – beak swabs (BS), blue – soil samples (So) and purple – talon swabs (TS). Shaded areas represent 95 % confidence intervals. Only faeces that was not potentially contaminated with *Salvelinus alpinus* was included. One eagle faeces containing only the Greenland halibut (*Reinhardtius hippoglossoides*) shared no dietary taxa with any other sample and was removed from the NMDS visualisation as the outlier compressed all remaining data points into a single line, making it impossible to visualise community structures.

**Figure S4:**
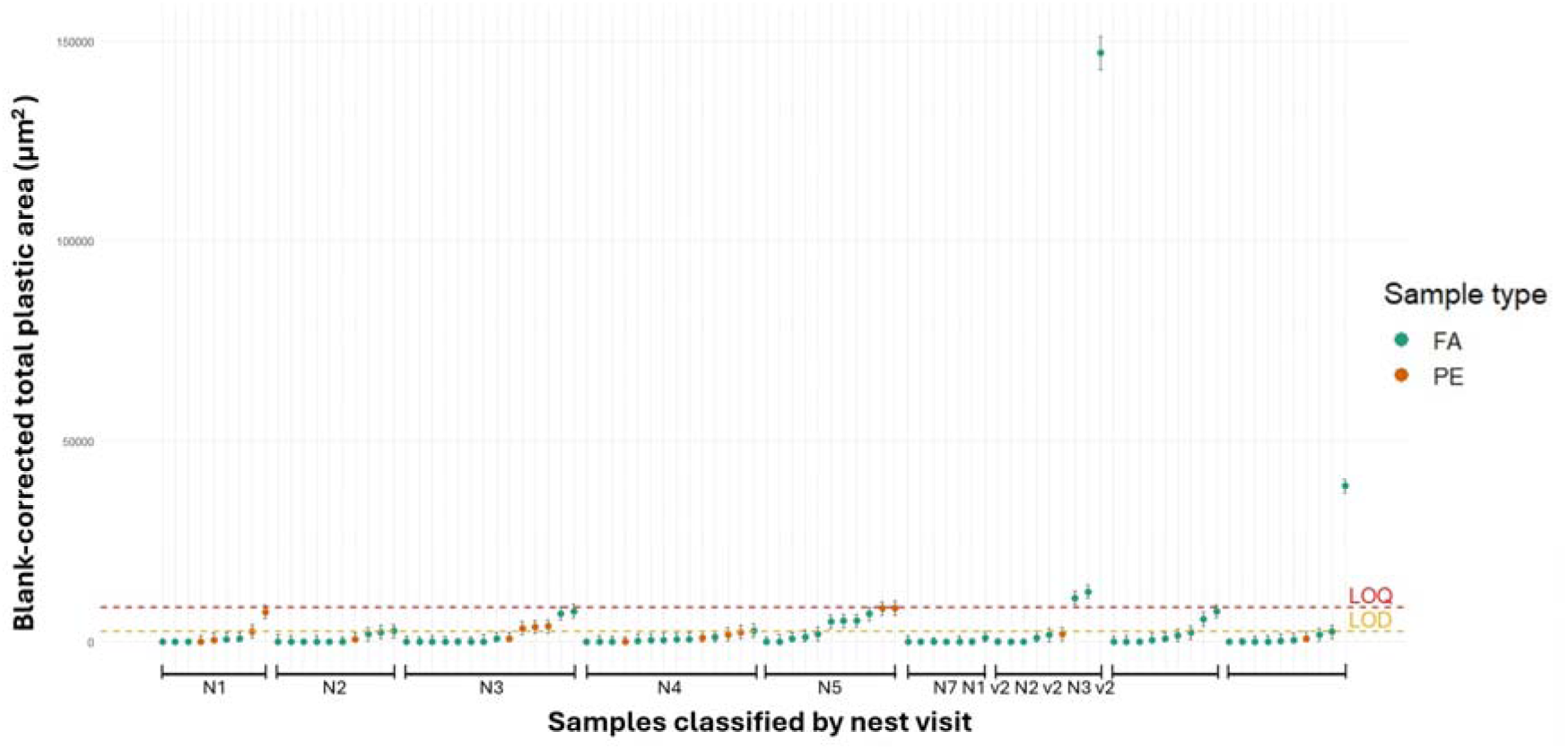
Total area of microplastic in pellets and faeces. Pellets (PE) are in orange and Faeces (FA) in green. Each data point is the best estimate of plastic area after blank correction using Monte Carlo uncertainty propagation. Bars show the 95 % Monte Carlo percentile intervals. Two thresholds are applied on the graph, the Limit of Quantification (LOQ), in red and the Limit of Detection (LOD), in orange. Samples above LOD are described as contaminated with microplastics and samples above LOQ can be described using quantifiable variables. The outlier (sample 181A) was taken in nest 1 and contained DNA from northern raven and Arctic char.

**Figure S5:**
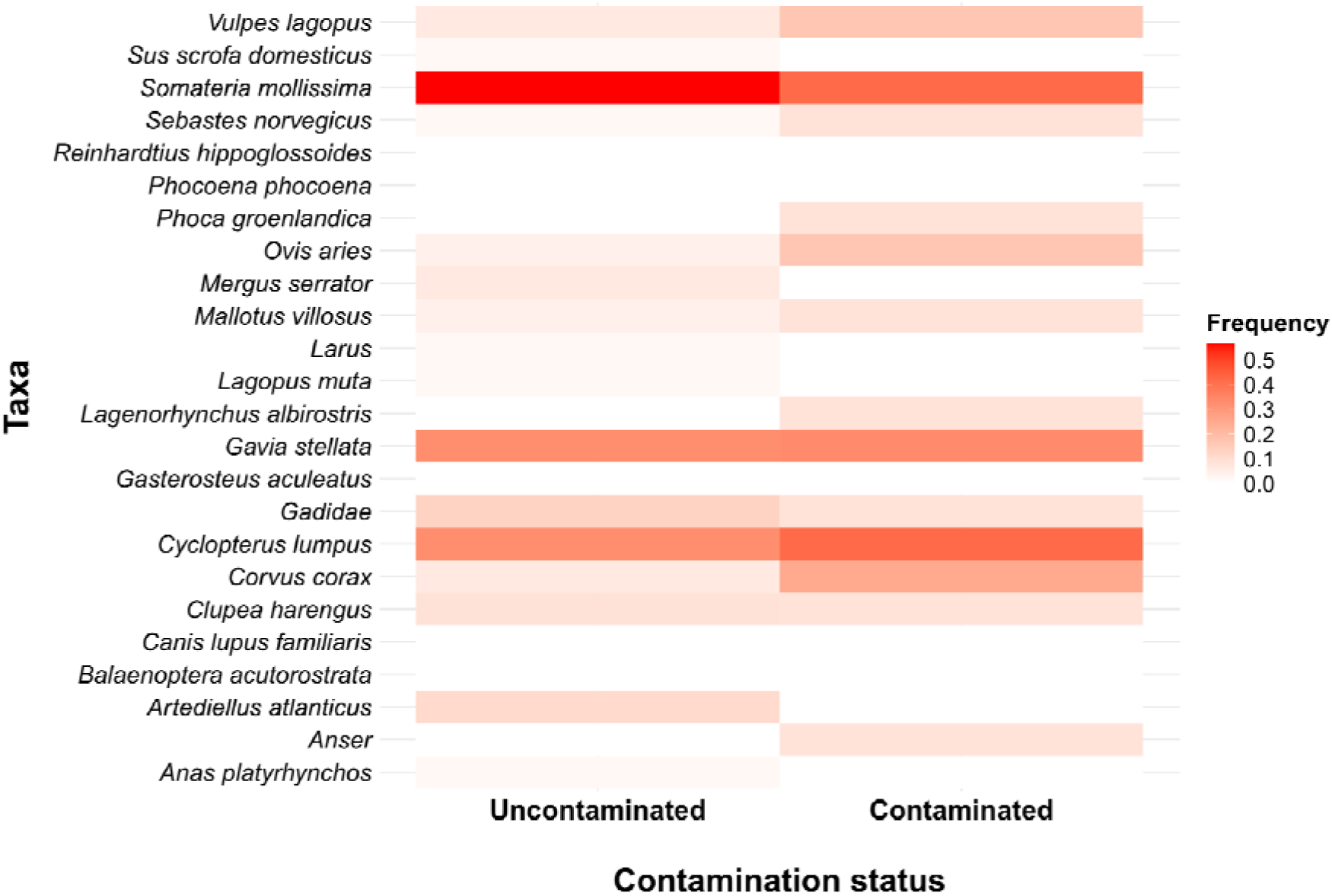
Heatmap comparing the frequency of dietary taxa present in faeces with and without plastic pollution, based on the detection thresholds. Only faecal samples for which both plastic assessment and diet analysis were done were included. Arctic char was removed from the analysis.

**Figure S6:**
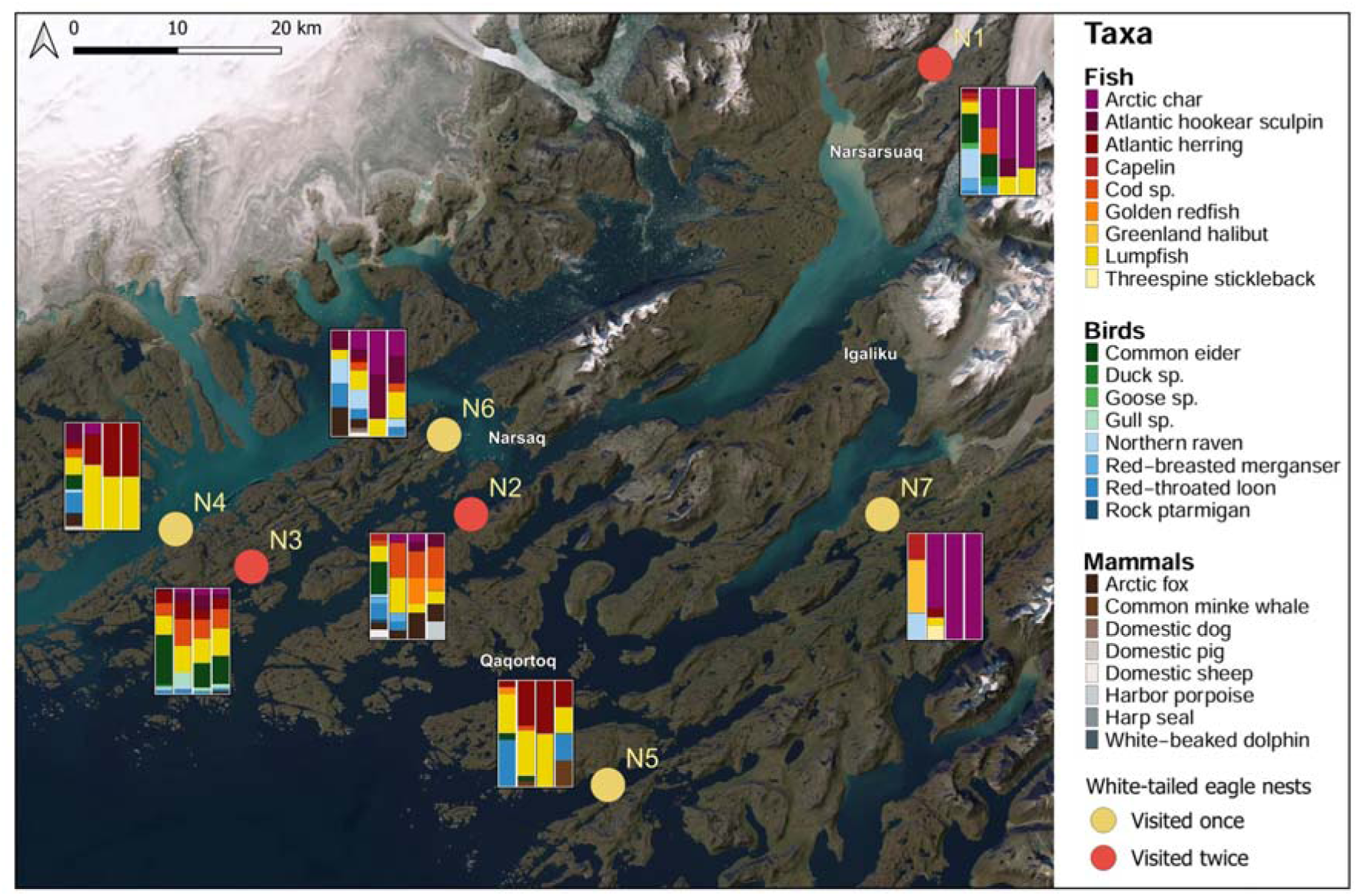
Map depicting the study area with each white-tailed eagle nesting sites and the diet obtained at each nest. Stacked bar plot at each nest displays weighted percent of occurrence (wPOO) data of prey items. From left to right, the columns show results of soil samples (So), beak swabs (BS), talon swabs (TS) and faeces (FA), respectively. Arctic char was excluded from faecal samples because of potential cross contamination. Nests in red were visited twice and nests in yellow were visited once.

### Supplementary tables

**Table S1:** Overview of the date of each nest visit with the number of chicks found, faeces collected and pellets collected.

| Nest ID | Date | Number of chicks swabbed | Number of faeces | Number of pellets |
| --- | --- | --- | --- | --- |
| 1 | 29/06/2025 | 1 | 5 | 5 |
| 2 | 04/07/2025 | 1 | 10 | 1 |
| 3 | 05/07/2025 | 3 | 10 | 4 |
| 4 | 05/07/2025 | 2 | 11 | 4 |
| 5 | 08/07/2025 | 1 | 10 | 2 |
| 2 (2<sup>nd</sup> visit) | 09/07/2025 | 1 | 10 | 0 |
| 6 | 11/07/2025 | 2 | 11 | 0 |
| 3 (2<sup>nd</sup> visit) | 12/07/2025 | 3 | 12 | 1 |
| 1 (2<sup>nd</sup> visit) | 14/07/2025 | 1 | 10 | 1 |
| 7 | 15/07/2025 | 1 | 10 | 0 |

**Table S2:** The total number of reads obtained per soil sample after removal of host DNA and possible contaminants.

| **White-tailed eagle** |  |
| --- | --- |
| Soil sample | Reads |
| 12A | 8065 |
| 13A | 27428 |
| 14A | 101701 |
| 15A | 12721 |
| 16A | 0 |
| 17A | 0 |
| 68A | 795727 |
| 69A | 0 |
| 70A | 18181 |
| 71A | 234840 |
| 72A | 0 |
| 73A | 0 |
| 74A | 91009 |
| 75A | 815194 |
| 76A | 693117 |
| 77A | 1413600 |
| 78A | 1359644 |
| 79A | 188801 |
| 80A | 937456 |
| 81A | 526781 |
| 82A | 1107611 |
| 87A | 847829 |
| 88A | 760844 |
| 89A | 1333963 |
| 90A | 367233 |
| 91A | 679475 |
| 92A | 1642312 |
| 93A | 1306107 |
| 94A | 1197615 |
| 187A | 1781168 |

